# Translocations spur population growth but exacerbate inbreeding in an imperiled species

**DOI:** 10.1101/2023.11.11.566550

**Authors:** Tyler Linderoth, Lauren Deaner, Nancy Chen, Reed Bowman, Raoul Boughton, Sarah W. Fitzpatrick

**Affiliations:** W.K. Kellogg Biological Station, Michigan State University, Hickory Corners, Michigan 49060; Flatwoods Consulting Group, Tampa, Florida 33610; Department of Biology, University of Rochester, Rochester, New York 14620; Archbold Biological Station, Venus, Florida 33960; Mosaic Fertilizer LLC, Tampa, Florida 33547; Department of Integrative Biology, Michigan State University, East Lansing, Michigan 48824; Ecology, Evolution, and Behavior Program, Michigan State University, East Lansing, Michigan 48824

**Keywords:** conservation, genomics, pedigree, Florida Scrub-Jay, fitness

## Abstract

Land and natural resource usage that supports human society can pose a risk to the survival of other species, spurring biodiversity loss. In extreme cases, when development threatens the existence of individuals, wildlife managers may perform mitigation translocations, relocating individuals out of harm’s way. We investigated the efficacy of mitigation translocations as a conservation strategy in Federally Threatened Florida Scrub-Jays using a dataset that provided unprecedented resolution into both the demographic and genomic outcomes of translocations. Over the course of seven years, a total of fourteen groups (51 jays) from five subpopulations that had been declining from agriculture and lack-of-fire driven habitat degradation were translocated to a larger site of more contiguous restored habitat with only four family groups, to mitigate for loss of these subpopulations from mining activity. Habitat restoration and translocations established a core population that increased 10-fold in size after only 17 years from the first translocations. Pedigree analyses of this population revealed that a small subset of mostly translocated individuals fueled the demographic expansion, with a single breeding pair responsible for ∼24% of the ancestral genetic contributions since 2008. Genomic comparisons between translocated individuals and individuals from the core population before and after translocations revealed that the high reproductive skew led to increased inbreeding and loss of genetic diversity. This study stresses the importance of demographic and genetic monitoring following translocations, and that subsequent, genetic-rescue-oriented translocations may be necessary in mitigation scenarios to counter the genetic consequences of reproductive skew in fragmented populations.

**Significance Statement:** There is ongoing debate surrounding the effectiveness of mitigation-driven translocations for conservation, however we show that translocations to mitigate the effects of mining on Federally Threatened Florida Scrub-Jays spurred population growth; a major boon to the viability of this species. We translocated individuals from at-risk subpopulations that were demographic sinks into recently restored habitat, which quickly established a rapidly growing core population. We demonstrate that demographic and genetic recovery do not necessarily go hand-in-hand, as reproduction was highly skewed towards a small subset of mostly translocated individuals, which increased inbreeding and eroded genetic diversity. This stresses the importance of demographic and genetic monitoring for identifying reproductive skew, allowing for adaptive management that addresses inbreeding and achieves broader conservation goals.

## Introduction

Human use of land and natural resources is a major driver of the modern biodiversity loss crisis (1), and its inevitable ongoing practice makes innovative and well-designed conservation strategies vital. Translocation, defined as human-mediated movement of live organisms (including gametes and propagules) from one area with release into another (2), is one conservation approach that can be particularly effective when habitat fragmentation restricts natural migration (3, 4). When translocations are used to reduce the detrimental impact on organisms resulting from human development they are classified as mitigation-driven translocations (5). Though mitigation translocations are primarily intended for relocating individuals that would otherwise be threatened with death, they should also improve the conservation status of the focal species or restore ecosystem functions as stated in the IUCN Guidelines for Reintroductions and Other Conservation Translocations (2), thus serving as a tool to combat biodiversity loss.

However, mitigation-driven translocation as a conservation management strategy is controversial, with the major critiques focused on low success rates, unscientifically sound execution, and lack of subsequent monitoring (5, 6). Often, the rapid time frame of human development projects necessitates that management action be taken prior to the extensive effort required to obtain and analyze sufficient biological data to make sound ecological and conservation risk assessments. Yet, in the face of slated (or already completed) habitat alteration, relocating individuals may be the only option to preserve lineages and ultimately conserve populations. One possible advantage of relocating individuals from multiple distinct populations, especially those that have become recently small and isolated, is the potential for restored gene flow and increased genetic variation that could result in genetic rescue (7). Understanding and being able to predict both the demographic and genetic outcomes in these scenarios is invaluable for implementing translocations that achieve broader conservation goals (8–12).

Here, for the first time, we show clear mechanistic links between the demographic and genomic consequences of mitigation translocations using a full population pedigree and temporal whole-genome sequencing data collected over two decades of detailed monitoring of a Florida Scrub-Jay (*Aphelocoma coerulescens*, hereafter FSJ) metapopulation. The Florida Scrub-Jay is a nonmigratory, endemic bird species to Florida, USA, that has experienced massive population decline starting in the twentieth century, mainly caused by degradation and loss of xeric oak scrub habitat on which it is reliant (13, 14). In 1987 FSJs were assigned Threatened status by the US Fish and Wildlife service (USFWS). This study is focused on FSJ subpopulations contained within a metapopulation called Metapopulation 4 (M4) (15) located on Florida’s west coast in an area now partially owned and operated by Mosaic Fertilizer LLC, which conducts phosphate mining activities. As part of a FSJ management plan developed between IMC Phosphates Company (now Mosaic Fertilizer, LLC) and the USFWS it was determined that multiple subpopulations on the periphery of the M4 metapopulation were demographic sinks residing in areas targeted for mining where the habitat had been degraded and fragmented from agriculture and lack of fire, and that translocating these birds into a larger site where habitat was more contiguous and had recently been restored, termed the Mosaic Wellfield (MW) site (Fig. 1A and 1B), could maximize the viability of the M4 metapopulation. From 2003 through 2010, jays from five M4 subpopulations facing high extinction risk (Fig. 1A) were annually translocated (except for 2006) into the MW site (Fig. 1C), totaling 51 translocated individuals. Prior to translocations and up through the present, all birds in the MW and neighboring protected sites (Duette Preserve and Coker Tract), collectively called the M4 Core Region (CR), were monitored, yielding extensive demographic data, a nearly complete population pedigree, and material for genetic analyses, providing an opportunity to fully characterize and understand in unprecedented detail the outcomes of mitigation translocations in a cooperative-breeding species of high conservation concern.

**Figure 1.**
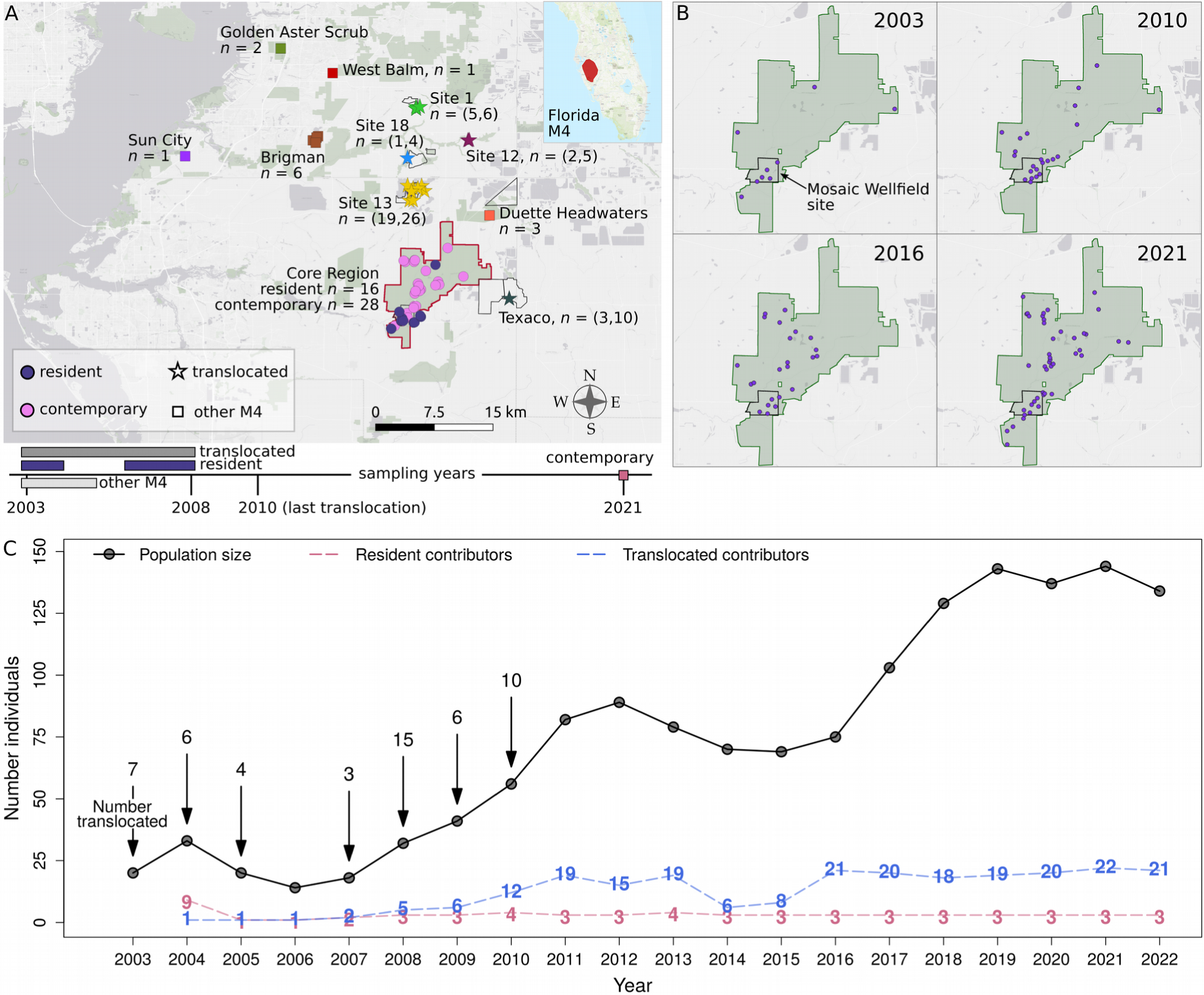
Demographic response of the Core Region FSJ population to translocations. (**A**) Map showing M4 subpopulation sample sizes (*n*) for genetic analyses, with individuals depicted by colored shapes. Sample size value pairs for translocation donor subpopulations denote (genetic sample size, total number individuals translocated from site). Individuals were translocated into the Mosaic Wellfield site and then expanded throughout the Core Region (CR) outlined in red. The geographic range of the entire M4 metapopulation is colored red in the inset map of the Florida peninsula. The time periods in which blood samples were taken from individuals for genetic analyses are displayed below the map. (**B**) Geographic expansion of family groups, denoted by purple dots, throughout the CR (green area) over time. The MW site is outlined in black. (**C**) Census size of the CR population over time (solid black line) with arrows showing the year and number of jays translocated into the MW site. Dashed lines indicate the number of translocated and resident ancestral lineages contributing to cohorts born in the CR since 2004.

## Results

### Translocations fuel demographic recovery

Thirteen individuals comprising four family groups resided in the M4 Core Region (CR) prior to translocations. These individuals along with three individuals born in the CR in 2004, 2005, and 2008 before the translocation period ended were considered CR residents (*n* = 16). The first four translocations had limited settlement success, so methodologies were adapted to better mimic natural FSJ dispersal patterns (16), resulting in improved settlement for translocations conducted in 2008–2010. Following each of the modified translocations, the CR population began to increase in size (Fig. 1C), while also expanding geographically as new family groups established territories throughout the CR (Fig. 1B). The CR population continued to grow rapidly years after the last jays were translocated, such that by 2022 the population had increased 10-fold, showing a clear positive demographic response to translocations. We refer to the CR population in years 2021 and on as the “contemporary” population.

### Population pedigree reveals reproductive skew

Out of 74 individuals that founded the CR population (resident and translocated jays), an increasingly small subset were genetically represented (either through themselves or their offspring) in the CR population over time (Fig. 2A and 2B), with translocated jays dominating the genetic contribution to cohorts born in the CR since 2008, halfway through the translocation period (Fig. 1C, Fig. S1). Since 2015, over 76% of the contemporary population’s founding ancestry, i.e. the genetic material originating from the founding individuals, is expected to have derived from the same 16 ancestors based on the population pedigree (Fig. 2A and 2B), which increased to 88% by 2022. Of these 16 ancestors, only three were resident jays.

**Figure 2.**
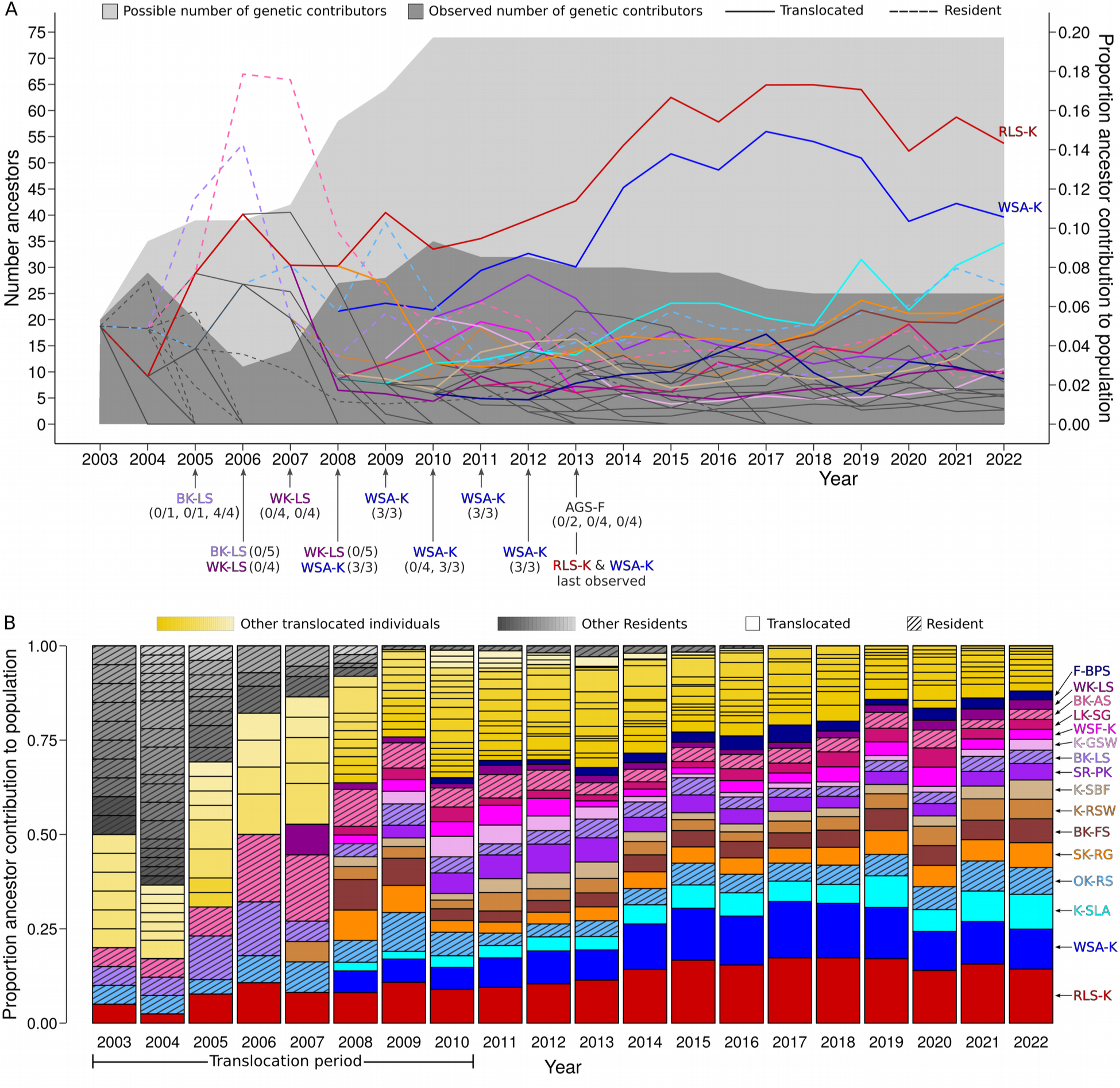
Expected genetic contributions by CR ancestors over time based on the population pedigree. (**A**) The total number of founding ancestors that had existed in the CR and could potentially contribute to the population is indicated by the top of the light gray shaded area, while the number of ancestors that were actually genetically represented in the population based on the pedigree is denoted by the top of the dark gray shaded area. The number of potential genetic contributors increased as jays were moved to the MW during the translocation period (2003–2010), with only a fraction contributing to the population over time. Lines show the expected proportion of the total genetic material originating from all ancestors of the CR population for each of the ancestors over time. Genetic contributions were skewed towards a small subset of CR founders, with one breeding pair comprised of RLS-K (male, red) and WSA-K (female, blue) dominating the reproductive skew. Annotated arrows show the mate of RLS-K each year and the number of young that the respective pair fledged out of the total number of eggs laid per nest attempt; (number fledged / total number eggs). (**B**) The same expected genetic contribution by ancestors to the CR population depicted by lines in (A), but shown as stacked bars, highlighting genetic takeover by a subset of mostly translocated lineages. Sixteen founding individuals from which over 76% of the CR ancestry is expected to have derived since 2015 are labeled in (B) and denoted by the same colors in (A). The remaining ancestral lineages are colored yellow (B) or gray (lines in A, bars in B).

The nearly complete population pedigree revealed that one breeding pair in particular, comprised of the male RLS-K and female WSA-K, made exceptionally high genetic contributions to the CR population (Fig. 2A and 2B). RLS-K was translocated into the MW from Site 13 in 2003 and after multiple seasons of low fledging success with two different females, paired with WSA-K, who had been translocated from Site 1 in 2008. This pair had high reproductive success every year from their first nest attempt in 2008 up until 2013 when they were last observed, fledging all offspring in 5/6 nest attempts, totaling 15 fledged offspring. On average, they collectively accounted for 24% (SD = 0.06) of the expected CR population founder ancestry, with a maximum contribution of 32% in 2017, four years after their presumed deaths indicating successful reproduction of their offspring. Since 2008, the average individual genetic contribution by RLS-K and WSA-K was respectively 2.4x and 1.8x higher than that of the next highest ancestor’s contribution to the CR population over this period. While this pair has dominated the reproductive skew, the contribution of K-SLA, another translocated male from Site 13 and relative of RLS-K (pedigree *r* = 0.125), has steadily increased, and by 2022 had nearly reached similar levels as WSA-K (Fig. 2A and 2B).

### Population genomic consequences of reproductive skew

Increased risk for inbreeding is a potential outcome of reproductive skew that can be particularly pronounced in small, isolated populations. We assessed the genetic consequences of skew in the CR population by sequencing the genome of one nestling from each 2021 CR breeding pair (contemporary sample, *n* = 28), which we compared to the genomes of 29 translocated jays and 16 residents likely from year classes 2002– 2008. We also sequenced the genomes of 13 birds from five other M4 subpopulations (Fig. 1A) and identified 2,049,176 high-quality SNPs segregating across all individuals. The donor populations were collectively genetically separated from the resident CR population by an *F_ST_* of 3%. In the space defined by the first two principal components (PCs) of a PC analysis (PCA) of genetic variation across space and time, contemporary individuals gravitated towards translocated individuals with high reproductive skew (Fig. 3A), suggesting that they are related to these highly reproductively successful individuals, while the overall contemporary CR population was 2.6% divergent from the resident population in terms of *F_ST_*. The majority of contemporary individuals in the PCA diffuse towards WSA-K, reflecting this individual’s large, multi-generational reproductive contribution (note that their mate, RLS-K, was not sequenced).

**Figure 3.**
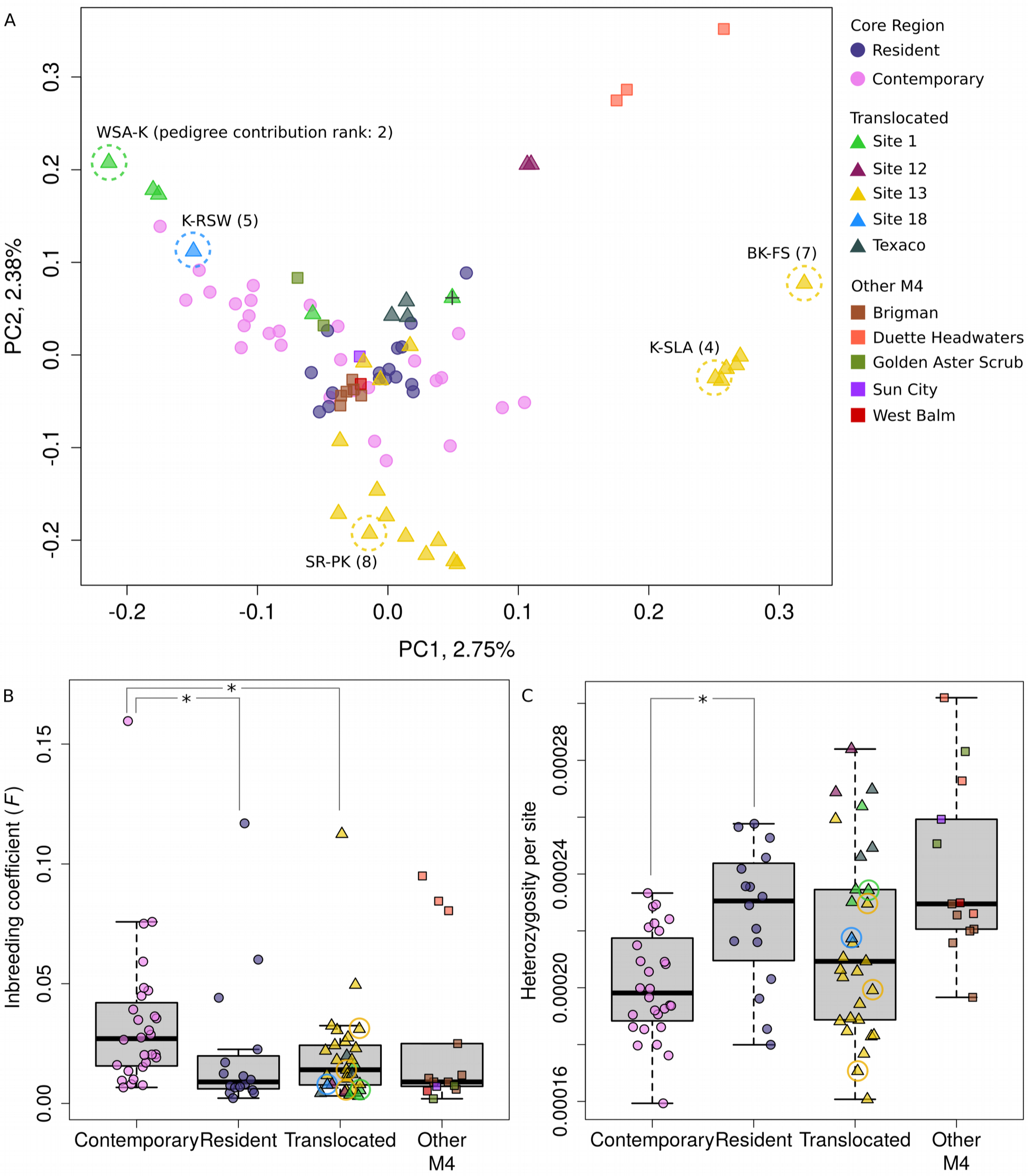
Spatio-temporal genomic characterization of the Core Region and neighboring M4 subpopulations. (**A**) Principal component analyses of the genome-wide variation among resident (historic) and contemporary CR individuals, translocated jays, and jays from peripheral M4 subpopulations. The percentage of the total genetic variation explained by each principal component (PC) is denoted on each axis. Sequenced ancestral individuals among the top 10 largest genetic contributors to the contemporary sample based on the pedigree are enclosed in dashed circles and denoted by their color-band identifiers and pedigree-based contribution rank value. ‘+’ denotes a donor site individual that was not translocated and not a member of the CR population. (**B** and **C**) Distributions of individual inbreeding coefficients (*F*) and heterozygosity (*H*) within the resident and contemporary CR population and among translocated jays and peripheral M4 subpopulation individuals. Colors and shapes denoting individuals are the same as in (A). Significant (*p*-value < 0.05) group differences in mean *F* and *H* are denoted by ‘*’. The same highest-ranked contributors to the contemporary CR population based on the pedigree indicated in (A) are also circled in the plots of *F* and *H*.

In line with a partial genomic sweep from reproductive skew, on average, inbreeding estimated from genomic data was significantly higher (Mann Whitney *U* = 114, two-tailed *p*-value = 0.007) among contemporary individuals (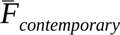 = 0.034, SD = 0.031) compared to residents (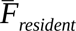 = 0.021, SD = 0.030) (Fig. 3B), while heterozygosity was significantly lower (Mann Whitney *U* = 350, two-tailed *p*-value = 0.002) among contemporary individuals (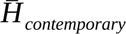 = 2.01 × 10^−4^, SD = 1.85 × 10^−5^) compared to residents (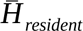 = 2.25 × 10^−4^, SD = 2.44 × 10^−5^) (Fig. 3C). Average inbreeding was also significantly higher (Mann Whitney *U* = 574, two-tailed *p*-value = 0.007) among contemporary individuals than among translocated jays (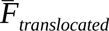 = 0.020, SD = 0.021), while levels of inbreeding were the same between resident and translocated individuals (Mann Whitney *U* = 199, two-tailed *p*-value = 0.445), suggesting that the increase in contemporary inbreeding is not accounted for by ancestral levels, and is the result of reproductive skew among founders. The contemporary population also had lower nucleotide diversity (average number of pairwise genetic differences), *π*, than the resident population (*π_contemporary_* = 2.36 × 10^−4^, *π_resident_* = 2.49 × 10^−4^) consistent with less heterozygosity, as well as lower genetic diversity based on the number of segregating sites as measured by Watterson’s *Θ*, *Θ_W_*, (*Θ_W,contemporary_* = 4.16 × 10^−4^, *Θ_W,resident_* = 4.28 × 10^−4^).

### Detecting skew using genetic data

We investigated approaches for characterizing reproductive skew solely from genetic data in order to 1) corroborate skew identified from the pedigree and 2) leverage the rare opportunity of having a real-life scenario involving a nearly complete population pedigree and genomic data to validate ways of identifying skew in the absence of a pedigree, which is the case for many population monitoring scenarios. We quantified the genetic contribution of sequenced ancestors to the contemporary population based on their average rank-weighted relatedness to all sequenced individuals from 2021, denoted *K* (see methods section Characterizing skew from genomic data).

The *K* statistic values were highly significantly correlated with pedigree-based estimates of the expected genomic representation of sequenced ancestors in the contemporary population (Kendall’s *τ* = 0.42, *z* = 3.35, *p*-value = 8.21 × 10^−4^). The distribution of *K* (Fig. 4A) is skewed towards the same individuals with disproportionately high contribution in the pedigree (Fig. 4A and 4B), and in agreement with the pedigree, WSA-K stands out as the topmost genetic contributor (since her highly successful mate, RLS-K, was not sequenced). Half of the top 10 individuals with the highest pedigree skew are also among the top 10 individuals with the highest *K* (Fig. 4B), and the remaining half were not sequenced, suggesting high power to identify relatively prolific lineages from genetic data alone. The *K* value for WSA-K was nearly twice as high as that for any other ancestor except for K-RSW, making WSA-K and K-RSW stand out in the distribution of *K* (Fig. 4A), indicating that it can be used to identify exceptionally high genetic contributors even against a background of reproductive skew. *K* for some individuals will be elevated by proxy of high relatedness to an individual with large reproductive contribution. Such is the case for GK-YS which is ranked as the third highest genetic contributor based on *K* despite making no reproductive contribution in the pedigree (Fig. 4A), which derives from very strong (*r* = 0.59) relatedness to WSA-K.

**Figure 4.**
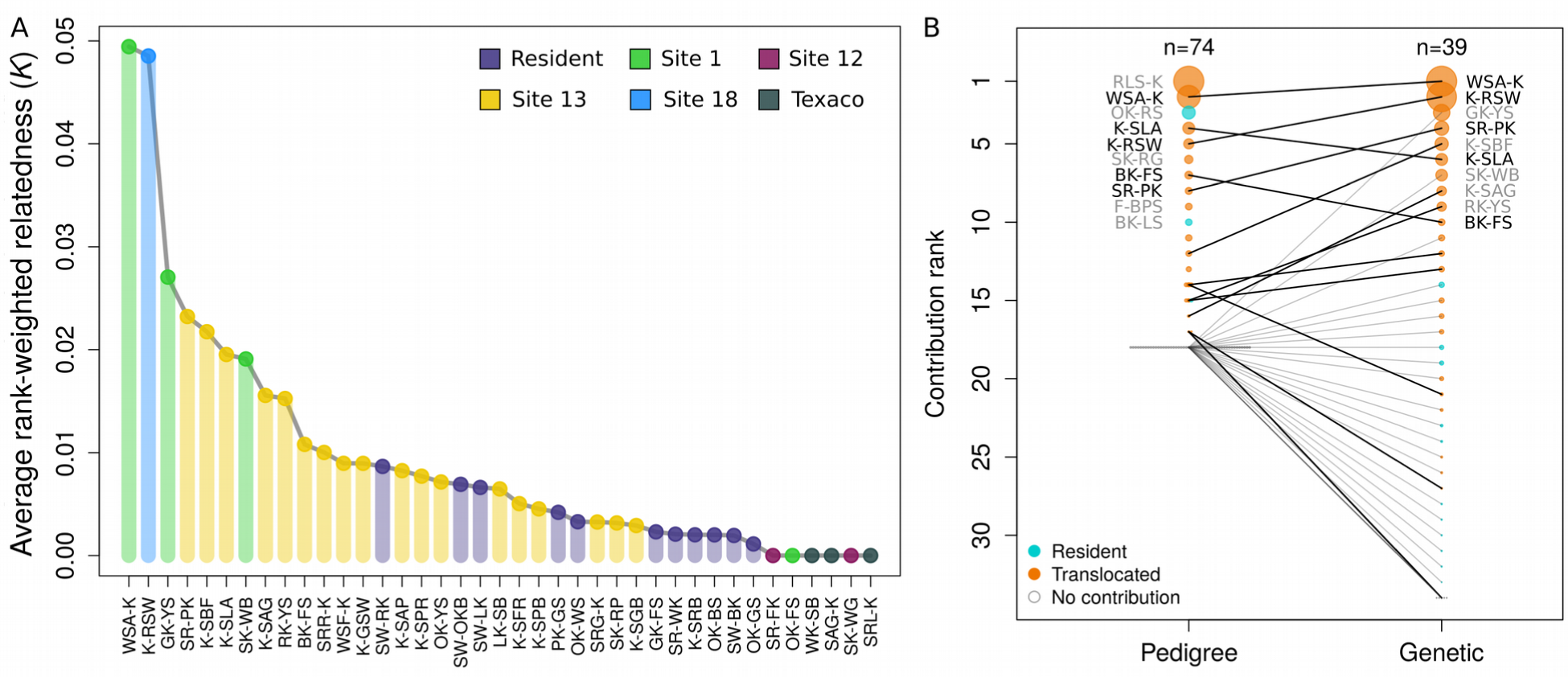
Characterization of reproductive skew from genomic sequencing data. (**A**) Distribution of the average rank-weighted relatedness statistic, *K*, for sequenced ancestral individuals, highlighting large differences in the genetic contribution among founders of the contemporary population. (**B**) Rank comparison of the genetic contribution to the contemporary CR population between the pedigree-based analysis involving all 74 ancestors and the genetic *K* statistic calculated for 39 sequenced ancestors, showing high concordance for individuals with high reproductive skew. Resident versus translocated origin for individuals is indicated by cyan and orange colors, respectively, and the size of points are scaled to the maximum individual contribution for the respective analysis. Lines connect individuals common to both analyses and are bold if the individual contributed to the 2021 CR population according to the pedigree. The identity of the top 10 contributing jays from each analysis are displayed next to their respective points with individuals common to both lists in bold.

## Discussion

We documented a 10-fold increase of the M4 Core Region FSJ population size within only twenty years of receiving 51 translocated jays from five neighboring M4 subpopulations between 2003 and 2010. Prior modeling had shown that establishing a CR of habitat and translocating the subpopulations (14 breeding pairs to one location) was the optimal mitigation and should increase the overall population viability for the M4 in the face of mining activity (17, 18). The demographic growth of the CR population strongly suggests that translocations, even for mitigation, can be an effective conservation tool. Following translocations, the CR population expanded geographically, with new family groups establishing *de novo* territories throughout expanding areas of restored habitat, clearly demonstrating the vital role that habitat management played in promoting population growth. While it is tempting to assume that such demographic increase is likely to be accompanied by less inbreeding and lower loss of genetic diversity, and perhaps even increased diversity when fueled by translocations, our genomic results show that this is not necessarily the case. In fact, we observed the opposite, whereby reproductive skew among founding lineages of the expanding population resulted in higher inbreeding and lower heterozygosity compared to levels in the resident CR population. Disproportionately high reproductive contribution tended to be from translocated individuals; in particular one breeding pair and their descendants have dominated genetic contributions to the CR population for the past 14 years.

Extreme reproductive skew can lead to inbreeding depression and even population collapse as most clearly demonstrated by the wolves of Isle Royale (19–21). While the partial genomic sweep of the RLS-K and WSA-K lineages throughout the CR population is far less severe, even slightly inbred FSJs (*F* = 0.008) have been shown to have lower rates of survival to yearling and breeding stages compared to entirely outbred individuals (22). And though reproductive skew is a common feature in many natural populations spanning diverse taxa, it may have especially dire consequences within recently small, fragmented populations experiencing decreased immigration. Increased inbreeding and loss of genetic diversity from reproductive skew have been documented in other translocation cases involving birds and other taxa (12, 23), suggesting that this is an issue that should be assessed in monitoring programs for populations augmented with or established via translocations. Realistically however, in many translocation scenarios it will be infeasible to carry out the type of population-level monitoring required to construct full pedigrees used to characterize skew analogously to this study. Fortunately, we demonstrate that skew can be reliably detected and quantified from molecular data alone using the average rank-weighted relatedness statistic, *K*, which showed strong congruence with pedigree-based inference in terms of which lineages had disproportionately high representation in the CR population. Identifying skew using *K* is possible whenever population genetic data from at least two time points permits calculating relatedness between ancestors and decedents. The ability for identifying exact individuals driving skew depends on their inclusion in the genetic sample (they will always have the highest *K*), however, if those individuals are absent in the sample, family or lineage-level resolution will still be possible through proxy of sampled individuals closely related to the actual skewed breeder, which will often be sufficient for determining the extent to which skew is occurring in a population. Therefore, for the purpose of characterizing skew using *K,* we recommend genetic sampling from as many ancestral resident or translocated individuals as possible and a representative population sample of potentially minimally related descendants appropriate for more general population genetic characterizations (e.g. estimating allele frequencies, inbreeding, etc.), which aligns well with conventional sampling schemes for population genetic monitoring.

Although the CR population looks to be on a less viable genetic trajectory as a result of mitigation-driven translocations, it is important to emphasize the demographic recovery that these translocations produced. Larger census size provides a buffer against demographic and environmental stochasticity, critical for population persistence (24). Also, the contribution of the RLS-K/WSA-K lineages to the CR population appears to have stabilized after ∼2015 and is on average gradually decreasing among cohorts born in subsequent years, suggesting that the threat of a continued genomic sweep of these lineages may be subsiding. However, the contribution from other highly-represented translocated lineages, particularly K-SLA, appear to be increasing. Given the negative genetic consequences of skew in this small population and its isolation due to habitat fragmentation, future translocations for the purpose of genetic rescue may be warranted in order to strategically combat inbreeding and increase the effective population size to one where genetic rescue becomes increasingly unnecessary. Such translocations may necessitate the import of genetic material from outside of the M4 metapopulation and will require care to avoid genomic sweeps.

Based on the findings of this study, we advocate the need for continued monitoring and potential management even after apparent demographic growth following translocations, which should be factored into the management plans for many cooperatively-breeding birds and other species in which reproductive skew is not uncommon (25, 26). Both demographic and genetic health are necessary for population viability and it is through vigilant monitoring, integrative data collection, and habitat management efforts that this has been partially achieved in the CR while enabling identification of negative outcomes that managers can design informed-solutions for in order to ensure long-term success. Thus, despite the critiques of mitigation-driven translocations we believe that this case study provides an excellent model for how they can be carried out to achieve broader conservation goals.

## Materials and Methods

### Samples and sequencing

We conducted CR population censuses every spring before the breeding season and every July at the end of the nesting season. We identified breeding pairs and tracked their reproductive efforts through banding and field behavior observations. Florida Scrub-Jays are genetically monogamous regardless of social factors and environment (27), allowing for accurate assignment of parentage from behavioral observations (28). Thus there is strong concordance between pedigrees constructed from genetic and observational data in this species. Nests were located and monitored for each family group to determine clutch size, hatching success, and fledging success. Adult birds were banded beginning in 1999 and thereafter any adults were banded and classified as immigrants. We began monitoring nests in 2004 and through 2017 young were banded as independents primarily between June and September each year. From 2018 through 2021, we banded and collected blood samples from 12-day-old nestlings using 25 gauge needles. We collected blood from translocated birds at the time of capture using 27.5 gauge needles. Blood was taken from the brachial vein and stored in lysis buffer for genetic analysis.

We sequenced DNA from the blood of 87 individuals, encompassing 13 individuals from all four family groups present in the MW prior to translocations and 28 individuals born in the CR in 2021, 19 years after translocations began. Resident jays were sampled from 2003–2008. We assigned five individuals (K-WS, S-GK, PK-SL, RK-SB, WK-SL) sampled in the MW in 2004, 2006, 2007, and 2008 that were banded as adults to the resident CR group based on clear genetic affiliation to the resident population. We sampled 28 of the 51 translocated jays in 2003–2008, as well as one individual (K-SBP) from Site 13 that naturally immigrated to the CR between July 2009 and March 2010 which we treated as translocated in the pedigree and genetic analyses. In 2005, we also sampled one non-translocated individual (RSW-K) from Site 1 who migrated to Little Manatee State Park, which is outside of the CR, and so was not considered translocated nor part of the CR population in any analyses. The remaining 13 individuals were sampled in 2003–2005 from five non-donor M4 subpopulations north of the CR.

We extracted DNA from blood samples using Qiagen DNeasy Blood and Tissue kits per the manufacturer’s protocols. Whole genomes for each sample were sequenced by Novogene Corporation Inc. on the Illumina NovaSeq 6000 using 150 bp paired end sequencing. Novogene carried out basic quality control on the raw sequencing reads.

Reads containing adapters sequence, greater than 10% undetermined bases (N), and/or more than 50% bases with Phred-scaled quality score below six were filtered out. Following quality control the amount of data per individual ranged from ∼7.76–14.89 billion reads (median = 8.40 x 10^9^ reads).

### Pedigree analysis

The expected number of genomic copies originating from each resident and translocated individual among all extant CR individuals as well as within CR cohorts was calculated annually based on a nearly complete population pedigree using the ‘pedstat’ function of relateStats (https://github.com/tplinderoth/PopGenomicsTools). This function implements the algorithm from (29) for calculating expected genomic contributions from a pedigree and relies on an additive genetic relatedness matrix, which was inferred using the ‘makeA’ function of the nadiv R package (30). The expected number of genome copies, *n_i_,* originating from a focal ancestor *i* from time *t* in a descendant group (population or cohort) consisting of individuals *M_t_*_+ Δ_ from a later time point, 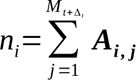. ***A_i, j_*** is a matrix consisting of relatedness coefficients, *r_i,j_*, between *i* and the *j*th individual of *M_t_*_+ Δ_ calculated from a pedigree for which links between individual *i* and their ancestors are removed. Cohorts were defined as all individuals born in the CR in a given year. Reproductive skew was inferred as disproportionately high expected genetic representation from a particular ancestral individual or group relative to the total genomic representation from all 74 founding ancestors.

### Genetic data quality control and mapping

Prior to mapping we performed additional quality control on the sequencing reads. First, we merged reads produced across different sequencing lanes for the same individual and then removed reads likely derived from PCR duplicates using SuperDeduper version 1.3.3 (31). We then used cutadapt version 3.7 (32) to trim residual adapters if at least the final three bases on the 3’ end of reads aligned to the start of the adapter sequences, trim missing bases (‘N’) from the ends of reads, and finally, remove pairs of reads for which at least one read was shorter than 70 bp. The clean reads for each individual were then separated according to sequencing lane so that the lane identity for different sequencing batches could be recorded in the alignment files.

We mapped quality-controlled reads to the FSJ reference genome assembly (33) using bwa mem version 0.7.17 (34) under default parameters. The ‘merge’ utility of SAMtools version 1.15.1 (35) was then used to combine aligned reads from different sequencing lanes into a single alignment (BAM) file per individual. SAMtools was used to add mate score tags to the reads with ‘fixmate’, sort the BAM files, and identify and record any residual PCR duplicates among the reads using ‘markdups’. The clipOverlap function of BamUtil version 1.0.15 (36) was used to clip overlapping sequence between paired reads from the mate with the lower average quality score. The median and mean sequencing depth per sample after quality-control and mapping was 6.9x and 7.2x (SD = 1.0), respectively.

### Genetic variant identification and quality control

We identified genetic variants using BCFtools version 1.15.1 (35) by first calculating individual genotype likelihoods (GLs) with the mpileup function from all uniquely mapped reads not derived from PCR duplicates and with a minimum Phred-scaled mapping quality of 20 and base quality of 13. We required at least two reads or 5% of the total reads be gapped within an individual for considering candidate indels. All other parameters were set to defaults for calculating genotype likelihoods. We called variants jointly across all 87 individuals using the BCFtools multiallelic ‘call -m’ model with the heterozygosity prior (-P) set to 0.003. Diploidy was assumed throughout except for Chromosome 24 (Z chromosome) for which we set females to haploid. For unsexed individuals (*n* = 53), we classified them as female if their Z chromosome to autosome average depth ratio was below 0.6 or male if this ratio was above 0.6 (see supporting information for further details). Sex assigned in this manner agreed with all 34 cases where sex had been assigned in the field based on behavioral observations or from an amplicon-based PCR assay.

We masked genomic sites deemed unreliable for downstream inference based on the distributions of sequencing and mapping quality metrics calculated from all autosomal and Z chromosome reads separately. Specifically, we masked out sites with excessively high or low total depth, low mapping quality, and low base quality across all samples. In addition, based on the subset of reads used for autosome and Z chromosome genotype calling respectively, we filtered out sites with excessively low and high high-quality-read depth (VCF DP flag) and excess heterozygosity across all individuals, as well as excessive missing data and/or low genotyping quality within the CR resident, CR contemporary, CR jays banded as adults, translocated, and all remaining M4 individual groups (see supporting information for quality masking details). All downstream population genetic analyses were limited to sites from 32 scaffolds homologous to zebra finch autosomes (we excluded the Z chromosome).

### Population genetic characterization

We calculated genotype likelihoods (GLs) for all high-quality, autosomal, biallelic SNPs with ANGSD version 0.937 (37) using the SAMtools model (-GL 1), considering only reads with minimum Phred-scaled map and base qualities of 20. We used the genotype likelihoods and prior probabilities for the genotypes under Hardy-Weinberg equilibrium given a maximum likelihood estimate of the sample-wide minor allele frequency (-doMaf 1) to obtain genotype posterior probabilities in ANGSD. The major and minor alleles for the allele frequency estimation were inferred from the genotype likelihoods (-doMajorMinor 1).

To assess spatio-temporal genetic structure we used the genotype posterior probabilities to calculate the genetic covariance among all individuals at SNPs with a minimum minor allele frequency (MAF) of 2% using ngsCovar, part of the ngsTools software package (38). We performed a PCA by decomposing the genetic covariance matrix in R version 4.1.3 (39) with the ‘eigen’ function. Pairwise differences in allele frequencies between spatial and temporal groups were quantified in terms of *F_ST_* in ANGSD using an estimation of the joint site frequency spectrum (SFS) between groups at all quality-controlled sites (see supporting information for details).

We measured genetic diversity of the resident and contemporary CR populations in terms of Watterson’s *Θ*, *Θ_W_*, and nucleotide diversity, *π*. Allele frequency likelihoods from ANGSD were used to calculate respective maximum likelihood estimates of the folded SFS for the resident and contemporary CR populations. Using prior probabilities for the MAF at any given site given by the folded SFS, we estimated posterior expectations of *Θ_W_* and *π* from the expected posterior estimate of the SFS with realSFS saf2theta. We also examined heterozygosity at the individual level, which we estimated based on the SFS for an individual in ANGSD (see supporting information). We also estimated individual inbreeding coefficients, *F*, from the genotype likelihoods at all high-quality SNPs with ngsF (40). For the expectation maximization algorithm of ngsF we used an initial estimate of F calculated from the GLs, and set the maximum number of iterations (--max_iters) to 3000, the minimum number of iterations (--min_iters) to 10, and the maximum root mean square deviation between iterations (--min_epsilon) to 1e-5 as the convergence criterion.

### Characterizing skew from genomic data

We used ngsRelate (41) to estimate relatedness between all pairs of individuals as the proportion of homologous alleles that are identical by descent (IBD) while accounting for the fact that individuals could be inbred as in (42). Specifically, the genotype likelihoods and allele frequency estimates from BCFtools at all quality-controlled SNPs were used to obtain maximum likelihood estimates for all nine condensed Jacquard coefficients that represent the possible IBD states between two diploid individuals, from which relatedness, *r*, was calculated. For every resident and translocated jay, we used these relatedness values to calculate the average rank-weighted relatedness to the contemporary sample, *K*. Specifically, for focal ancestor *i*, descendant individual *j*, and another randomly chosen ancestor 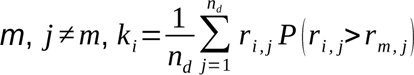, where *n_d_* is the number of descendant individuals and *r_i,j_* is the probability that alleles from *i* and *j* are IBD (i.e. the relatedness coefficient between *i* and *j*). Given *n_a_* ancestral individuals, 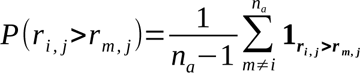. The *K* statistic will increase in value as the focal ancestor is more related to individuals in the descendant population, proportional to how much more they are related to the descendants than any other ancestors. Comparing values for *K* among ancestors is useful for identifying and assessing relative degrees of reproductive skew.

### Data and software availability

All data and code used to carry out analyses and generate figures for this study will be made publicly available upon publication of this manuscript. Computational tools developed in this study to calculate expected genetic contributions from pedigrees and the *K* statistic are implemented in the relateStats program at https://github.com/tplinderoth/PopGenomicsTools.

## Author Contributions

T.L., S.W.F., R. Boughton, R. Bowman, L.D., and N.C. designed and performed research and wrote the paper. L.D., R. Boughton and R. Bowman undertook field collections and data management. L.D., R. Boughton, R. Bowman, and T.L. performed data quality control. T.L. and L.D. analyzed data. T.L. contributed new analytical tools.

## Acknowledgments

We would like to posthumously thank Dr. Reed Bowman for his foresight and long term (>20 years) advising on the data collection and direction of the translocation of Florida Scrub-Jays in the M4 population. He made a big difference to the conservation of the population. We would also like to thank David Gordon who managed the initial project, banding and field data collections for 15 years as well as Lee Walton who also collected blood samples. The ongoing commitment and funds to support the extensive collection of field information has been supplied by Mosaic Fertilizer, LLC, and its predecessor IMC Phosphates. Support for this publication has also been provide by FSJ mitigation fees managed by the USFWS and distributed by The Nature Conservancy of Florida. S.W.F. is supported by NSF grant DEB 2016569. N.C. is supported by NIH grant R35 GM133412. This is KBS contribution 2370.

## Supporting Information Text

### Supporting Materials and Methods

#### Blood sample collection

Once a bird was captured, we rubbed the underwing with an alcohol swab and used a 27.5 gauge needle to lightly puncture the brachial vein. After removing the needle, capillary tubes were used to collect a blood sample. Each sample was transferred to a vial of lysis buffer labeled with the individual’s federal band number and the date of collection for storage. We used the same methods for samples collected prior to 2018, except that adults were captured with peanut-baited walk-in traps and we used 25 gauge needles for collecting blood.

#### Sex classification from next generation sequencing data

In order to assign sex to individuals without preexisting sex information (*n* = 53) we used SAMtools ‘coverage’ to calculate average autosome depth and the average Z chromosome depth, considering only uniquely mapped reads not sourced from PCR duplicates (-f ‘UNMAP,SECONDARY,QCFAIL,DUP’) and without imposing any maximum depth cutoff (-d 0) or minimum mapping quality or base quality thresholds (-q 0 -Q 0). Individuals with a Z chromosome to autosome average depth ratio below 0.6 were classified as females and individuals for which this ratio was greater than 0.6 as males since we expect the Z-chromosome depth to be approximately half that of the autosomes. Sex assigned in this manner agreed with all 34 cases where sex had been assigned in the field based on behavioral observations or from an amplicon-based PCR assay.

#### Genomic quality masks

We masked genomic sites deemed unreliable for downstream inference based on genome-wide distributions of sequencing and mapping quality metrics calculated separately from all autosomes and the Z chromosome. We masked out any sites with total depth (depth summed across all individuals) below the 0.5 quantile (523x for autosomal sites, 386x for Z chromosome sites) or total site depth above the 0.95 quantile (741x for autosomes, 681x for Z chromosome) or sites covered by zero reads. We also masked sites meeting at least one of the following criteria: 1) fraction of reads with map quality of zero greater than 2x the genome-wide average (10%), 2) root mean square (RMS) map quality below 35, 3) fraction of reads with base quality of zero greater than 2x the genome-wide average (0.3%), 4) RMS base quality below 20. We also filtered sites using quality metrics based on distributions for the subset of reads used for the genotype calling. We masked sites with total depth (VCF DP flag) above or below the respective autosomal/Z chromosome-wide median site depths +/− 25% (< 474x or > 790x for autosomes, < 336x or > 560x for Z chromosome) and/or with an exact test p-value for excess heterozygosity below the 2 percentile of the autosomal/Z chromosome-wide distributions (0.06 for both autosomes and Z chromosome). We filtered for minimum representation from different populations or groups by requiring at least 83% of individuals from each of the CR resident, CR contemporary, CR jays banded as adults, translocated, and all remaining M4 individuals be covered by at least two high-quality reads (minimum map and base qualities of 20 and 13, respectively) and have called genotypes in order to retain the site for downstream analyses. For potentially variable sites we required that the genotype quality (GQ) be at least 15 in 83% of individuals from each of the five aforementioned groups, otherwise the site was masked out.

#### Population genetic characterization

In order to estimate expected joint site frequency spectra used for calculating *F_ST_*, we first estimated the likelihoods of all possible derived allele frequencies at each high-quality, biallelic SNP (-doSaf 1) from the genotyping likelihoods for resident CR individuals, contemporary CR individuals, and the collective group of all translocated jays. We assumed that the reference allele represented the ancestral state noting that mispolarization of the ancestral state does not matter for calculating *F_ST_*. We then used these per SNP allele frequency likelihoods to obtain a maximum likelihood estimate of the pairwise joint allele frequency spectrum between groups using the realSFS subprogram of ANGSD. Lastly, using the allele frequency likelihoods for each group and the prior probabilities for observing joint allele frequencies provided by the 2-dimensional SFS, we estimated the within and between group genetic variance components of Reynold’s *F_ST_* estimator with realSFS. ‘Weighted’ genome-wide *F_ST_* were calculated as the ratio of the sum of between group to the sum of within group genetic variance over all sites.

We estimated heterozygosity, *H*, of an individual based on their site frequency spectrum using ANGSD. Specifically, for each individual, we first estimated the likelihood of having 0, 1, or 2 alternate alleles (-doSaf 1 supplying the reference genome to -anc) from the genotype likelihoods at all high-quality sites. We then used realSFS with these allele frequency likelihoods to estimate the folded SFS for each individual, which provides a maximum likelihood estimate of heterozygosity as 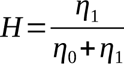, where *η*_1_ and *η*_0_ are the expected number of sites with zero and one minor alleles, respectively.

## Supporting Figures

**Fig. S1.**
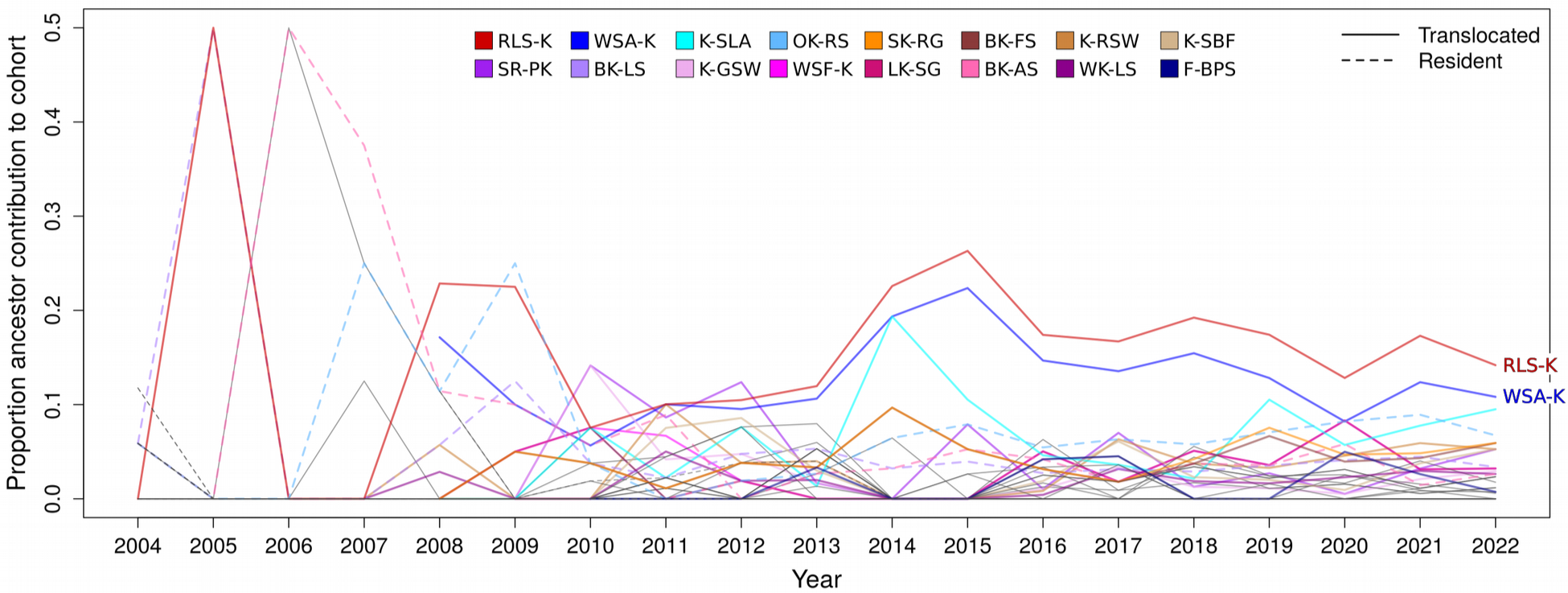
Expected genetic contribution to Core Region cohorts by ancestral individuals. Expected proportion of the genetic contribution by 74 ancestors to cohorts born in the CR from 2004–2022 based on the population pedigree. Sixteen lineages from which over 76% of the CR genetic ancestry originated since 2015 are colored. The breeding pair {RLS-K, WSA-K} made particularly high genetic contributions to CR cohorts since 2008.

## References

1. G. Ceballos, P. R. Ehrlich, Mammal population losses and the extinction crisis. Science 296, 904–907 (2002).

2. IUCN/SSC, “Guidelines for reintroductions and other conservation translocations” Version 1.0, (IUCN Species Survival Commission, 2013).

3. J. T. Hogg, S. H. Forbes, B. M. Steele, G. Luikart, Genetic rescue of an insular population of large mammals. Proc. R. Soc. B Biol. Sci. 273, 1491–1499 (2006).

4. W. E. Johnson et al., Genetic restoration of the Florida panther. Science 329, 1641–1645 (2010).

5. J. M. Germano et al., Mitigation-driven translocations: are we moving wildlife in the right direction? Front. Ecol. Environ. 13, 100–105 (2015).

6. H. S. Bradley, S. Tomlinson, M. D. Craig, A. T. Cross, P. W. Bateman, Mitigation translocation as a management tool. Conserv. Biol. 36, e13667 (2022).

7. A. R. Whiteley, S. W. Fitzpatrick, W. C. Funk, D. A. Tallmon, Genetic rescue to the rescue. Trends Ecol. Evol. 30, 42–49 (2015).

8. A. R. Weeks et al., Assessing the benefits and risks of translocations in changing environments: a genetic perspective: Translocations in changing environments. Evol. Appl. 4, 709–725 (2011).

9. E. S. Menges, Restoration demography and genetics of plants: when is a translocation successful? Aust. J. Bot. 56, 187–196 (2008).

10. S. Marić et al., Perils of brown trout (*Salmo* spp.) mitigation-driven translocations: a case study from the Vlasina Plateau, Southeast Serbia. Biol. Invasions 24, 999–1016 (2022).

11. L. D. Bertola et al., Genetic guidelines for translocations: Maintaining intraspecific diversity in the lion (*Panthera leo*). Evol. Appl. 15, 22–39 (2022).

12. K. A. Miller, N. J. Nelson, H. G. Smith, J. A. Moore, How do reproductive skew and founder group size affect genetic diversity in reintroduced populations? Mol. Ecol. 18, 3792–3802 (2009).

13. G. E. Woolfenden, J. W. Fitzpatrick, “Florida Scrub-Jay (*Aphelocoma coerulescens*)” in Birds of North America version 2.0, A. F. Poole, F. B. Gill, Eds. (Cornell Lab of Ornithology, 1996) 10.2173/bna.228 (accessed 7 August 7 2023).

14. J. W. Fitzpatrick, R. Bowman, “Florida scrub-jays: Oversized territories and group defense in a fire-maintained habitat” in Cooperative Breeding in Vertebrates 1st Ed., W. D. Koenig, J. L. Dickinson, Eds. (Cambridge University Press, 2016), pp. 77–96.

15. B. M. Stith, J. W. Fitzpatrick, G. E. Woolfenden, B. Pranty, “Classification and Conservation of Metapopulations; A Case Study of the Florida Scrub Jay” in Metapopulations and Wildlife Conservation, D. R. McCullough, Ed. (Island Press, 1996), pp. 187–215.

16. R. Bowman, L. Deaner, R. K. Boughton, Assessing conservation benefits of a successful mitigation translocation of Florida Scrub-jays, in prep.

17. U.S. Fish and Wildlife Service, “Scrub-Jay population modeling” in Biological Opinion and Associated Florida Scrub-Jay Habitat Management Plan for IMC Phosphates Company Southern Hillsborough and Manatee County Projects, (U.S. Fish and Wildlife Service, 2001), section 2.4 appendix 3.

18. R. Bowman, “Modelling the effect of different mitigation options for Florida Scrub-Jays (Aphelocoma coerulscens) on the Four Corners/Lonesome Regional Permit Area on the long-term persistence of the scrub-jay metapopulation in southern Hillsborough and Manatee Counties” (IMC-Agrico Company, 2008).

19. J. R. Adams, L. M. Vucetich, P. W. Hedrick, R. O. Peterson, J. A. Vucetich, Genomic sweep and potential genetic rescue during limiting environmental conditions in an isolated wolf population. Proc. R. Soc. B Biol. Sci. 278, 3336–3344 (2011).

20. J. A. Robinson et al., Genomic signatures of extensive inbreeding in Isle Royale wolves, a population on the threshold of extinction. Sci. Adv. 5, eaau0757 (2019).

21. P. W. Hedrick, J. A. Robinson, R. O. Peterson, J. A. Vucetich, Genetics and extinction and the example of Isle Royale wolves. Anim. Conserv. 22, 302–309 (2019).

22. N. Chen, E. J. Cosgrove, R. Bowman, J. W. Fitzpatrick, A. G. Clark, Genomic Consequences of Population Decline in the Endangered Florida Scrub-Jay. Curr. Biol. 26, 2974–2979 (2016).

23. I. G. Jamieson, Founder Effects, Inbreeding, and Loss of Genetic Diversity in Four Avian Reintroduction Programs: Inbreeding in Reintroduction Programs. Conserv. Biol. 25, 115–123 (2011).

24. V. Grimm, et al., Importance of Buffer Mechanisms for Population Viability Analysis. Conserv. Biol. 19, 578–580 (2005).

25. J. Haydock, W. D. Koenig, Patterns of Reproductive Skew in the Polygynandrous Acorn Woodpecker. Am. Nat. 162, 277–289 (2003).

26. N. J. Raihani, T. H. Clutton-Brock, Higher reproductive skew among birds than mammals in cooperatively breeding species. Biol. Lett. 6, 630–632 (2010).

27. A. K. Townsend, R. Bowman, J. W. Fitzpatrick, M. Dent, I. J. Lovette, Genetic monogamy across variable demographic landscapes in cooperatively breeding Florida scrub-jays. Behav. Ecol. 22, 464–470 (2011).

28. J. S. Quinn, G. E. Woolfenden, J. W. Fitzpatrick, B. N. White, Multi-locus DNA fingerprinting supports genetic monogamy in Florida scrub-jays. Behav. Ecol. Sociobiol. 45, 1–10 (1999).

29. D. C. Hunter, J. M. Pemberton, J. G. Pilkington, M. B. Morrissey, Pedigree-Based Estimation of Reproductive Value. J. Hered. 110, 433–444 (2019).

30. M. E. Wolak, nadiv: an R package to create relatedness matrices for estimating non-additive genetic variances in animal models. Methods Ecol. Evol. 3, 792–796 (2012).

31. K. R. Petersen, D. A. Streett, A. T. Gerritsen, S. S. Hunter, M. L. Settles, “Super deduper, fast PCR duplicate detection in fastq files” in Proceedings of the 6th ACM Conference on Bioinformatics, Computational Biology and Health Informatics, (ACM, 2015), pp. 491–492.

32. M. Martin, Cutadapt removes adapter sequences from high-throughput sequencing reads. EMBnet.journal 17, 10–12 (2011).

33. F. Romero et al., A new high quality genome assembly for the threatened Florida Scrub-Jay (Aphelocoma coerulescens), in prep.

34. H. Li, Aligning sequence reads, clone sequences and assembly contigs with BWA-MEM. arXiv [Preprint] (2013). 10.48550/arXiv.1303.3997 (accessed 12 April 2021).

35. P. Danecek et al., Twelve years of SAMtools and BCFtools. GigaScience 10, giab008 (2021).

36. G. Jun, M. K. Wing, G. R. Abecasis, H. M. Kang, An efficient and scalable analysis framework for variant extraction and refinement from population-scale DNA sequence data. Genome Res. 25, 918–925 (2015).

37. T. S. Korneliussen, A. Albrechtsen, R. Nielsen, ANGSD: Analysis of Next Generation Sequencing Data. BMC Bioinformatics 15, 356 (2014).

38. M. Fumagalli, F. G. Vieira, T. Linderoth, R. Nielsen, ngsTools: methods for population genetics analyses from next-generation sequencing data. Bioinforma. Oxf. Engl. 30, 1486–1487 (2014).

39. R Core Team, R: A Language and Environment for Statistical Computing. https://www.R-project.org/. (R Foundation for Statistical Computing, 2022).

40. F. G. Vieira, M. Fumagalli, A. Albrechtsen, R. Nielsen, Estimating inbreeding coefficients from NGS data: Impact on genotype calling and allele frequency estimation. Genome Res. 23, 1852–1861 (2013).

41. K. Hanghøj, I. Moltke, P. A. Andersen, A. Manica, T. S. Korneliussen, Fast and accurate relatedness estimation from high-throughput sequencing data in the presence of inbreeding. GigaScience 8 (2019).

42. P. W. Hedrick, R. C. Lacy, Measuring Relatedness between Inbred Individuals. J. Hered. 106, 20–25 (2015).

